# Information silos distort biomedical research

**DOI:** 10.1101/2021.07.26.453749

**Authors:** Raul Rodriguez-Esteban

## Abstract

Information silos have been an oft-maligned feature of scientific research for introducing a bias towards knowledge that is produced within a scientist’s own community. The vastness of the scientific literature has been commonly blamed for this phenomenon, despite recent improvements in information retrieval and text mining. Its actual negative impact on scientific progress, however, has never been quantified. This analysis attempts to do so by exploring its effects on biomedical discovery, particularly in the discovery of relations between diseases, genes and chemical compounds. Results indicate that the probability that two scientific facts will enable the discovery of a new fact depends on how far apart these two facts were published within the scientific landscape. In particular, the probability decreases exponentially with the citation distance. Thus, the direction of scientific progress is distorted based on the location in which each scientific fact is published, representing a path-dependent bias in which originally closely-located discoveries drive the sequence of future discoveries. To counter this bias, scientists should open the scope of their scientific work with modern computational approaches.

## 1. Introduction

The wide communication of scientific discoveries across the scientific community is an essential element of scientific research. Scientific silos have long been bemoaned for hindering this process (e.g. (Leischow et al., 2008; Vodovotz & An, 2013; Törmä, 2019)) by introducing a bias towards knowledge that is produced within a scientist’s own community. Analogous to corporate knowledge silos, there are at least three aspects that would define them: (1) enormous growth in the knowledge available to scientists, (2) organization of scientists into communities and (3) slowing of the propagation of scientific knowledge between those communities. Regarding the first aspect, the growth of information available for scientific research (Larsen PO & von Ins M, 2010; Bornmann & Mutz, 2015) represents a challenge for individual scientists as information seekers. In a perfect world, scientists would possess complete knowledge of all existing scientific information and select their research goals accordingly. Abundance of information, however, can represent its own “resource course” challenge. One could paraphrase the famous corporate knowledge-management adage (Sieloff, 1999) by saying: if only science knew what science knows. In this respect, the field of literature-based discovery (LBD) has propounded the existence of “undiscovered public knowledge” concerning facts that have never been put together because of the disparate venues in which they were published (Swanson, 1986; Bekhuis, 2006; Thilakaratne et al., 2019). Thus, there is a recognition that the milieu in which a discovery is published influences its later use by the scientific community due to the sheer abundance of existing scientific knowledge.

With respect to the second aspect, it has been shown that scientific publications are anchored around communities of scientists (Bruggeman et al., 2012; Shia et al., 2015; Fortunato et al., 2018), which go beyond traditional scientific communities (e.g. university departments, scientific organizations), representing a self-organizing process. This process might be encouraged by an institutional bias against interdisciplinary research (Bromhan & Dinnage, 2016; Baumwol et al., 2011), which would hamper collaboration across communities, despite recent trends towards fostering interdisciplinary research in systems and translational sciences (Luke et al., 2015; Auffray et al., 2009). It could also be a consequence of human cognitive limitations, due to scientists’ bounded capacity to learn and produce new knowledge and as a response to an increasingly more complex scientific landscape (Rodriguez-Esteban & Loging, 2013).

The third aspect, and the focus of this study, relates to the negative impact that scientific silos ultimately have on scientific progress. The existence of silos would entail that intra-silo information exchange is more frequent and faster than inter-silo. This would increase the likelihood of certain discoveries based on facts published within a silo, to the detriment of discoveries based on facts coming from different silos. Because new discoveries feed on past discoveries in a path-dependent manner (Soler et al., 2015; Tambolo, 2017; Heimeriks & Boschma, 2014), this dynamic could affect the long-term outcome of scientific research.

## 2. Research Objective

While siloization, and solutions that try to address it, have been a recurrent topic of scientific debate, no effort has been made to-date to quantify its negative impact on scientific progress, particularly its effect on the slowdown in the propagation of scientific facts, leading to the delay of certain discoveries and to the acceleration of others. This first attempt focuses on measuring the propagation of scientific facts about relations between compounds, genes and diseases, which are of broad interest in biomedical discovery, including clinical, pharmaceutical and translational research. Because defining silos is challenging, a surrogate distance measure—the citation distance—is used to represent the separation between publications within the scientific landscape. This measure would be related to the likelihood that publications belong to the same silo. Results of the analysis show that the citation distance between two published facts influences the probability that they will lead to a new discovery and thus signal the importance that knowledge silos (or, more broadly, the large-scale structure of relations between scientific publications) have in distorting scientific progress.

## 3. Methods

Scientific discovery can be modeled as a process in which facts are progressively connected to each other, thereby building growing networks in which the discovery of new facts is connected to already discovered facts (Cokol et al., 2005; Rzhetsky et al., 2015). The scientific discovery model employed in this study is inspired by the ABC model used in literature-based discovery (LBD) (Smalheiser, 2012; Thilakaratne et al., 2019) and it is based on undirected networks of up to 3 nodes (A, B and C). The nodes are particular elements that are the focus of research and the edges are relations between those elements that have been published in scientific publications. These networks are built sequentially over time: the edge AB is associated to the relation that is published first, the edge BC is associated to the second one, and the edge AC to the third one. Based on the time sequence order, the nodes are labeled appropriately as A, B or C. At any given point in time, and based on the existing published literature, there are networks with 1, 2 and 3 edges. For networks with all 3 edges, we say that AB and BC *enabled* the discovery of AC, even if there is no direct evidence of that, by virtue of precedence. AB and BC are considered “enabling facts” and AC, a “new discovery.” Networks with 2 edges comprise *potentially* enabling pairs of facts (i.e. AB and BC), which could enable a new discovery AC in the future.

In a full, three-edge network, the time lapse for a new discovery is the time between the publication of BC and the publication of AC. In a two-edge network (i.e. AC does not exist), the time lapse is measured between publication of BC and the cut-off time (January 1, 2020). This is done because potentially enabling facts can still enable a new discovery at a future date. This is handled analogously to a Kaplan-Meier curve to avoid biases due to right-censoring. One-edge networks are not considered for this calculation.

In this study, each network node (A, B, C) is one of each a gene, a disease or a compound. Each edge is a relation (e.g. a gene-disease relation) linked to a specific publication in the database MEDLINE. Data about relations came from The Comparative Toxicogenomics Database (CTD) (Davis et al., 2019), which was downloaded on May 4th, 2020. From this database, 1,603,976 unique relations between chemicals and genes were extracted; 34,830 relating genes and diseases and 218,868 relating chemicals and diseases. Additionally, cooccurrence data came from the MeSH and gene2pubmed databases. Chemical/drug and disease annotations were MeSH term annotations designated as “Major Topic” from the “Chemicals and Drugs” (D) and “Diseases” (C) branches, respectively, in the 2020 MeSH tree. Gene annotations came from the gene2pubmed database (Maglott et al., 2011) downloaded on August 20, 2020. These comprised 1,515,080 human gene annotations from 664,085 MEDLINE articles. MEDLINE data came from the 2020 MEDLINE/PubMed baseline. The reference date for each publication was the publication date (PubDate).

The citation distance was computed as the distance between nodes in an undirected citation network in which the nodes were scientific publications recorded in MEDLINE and connections were citations between them (Rodriguez-Esteban R, 2020; Rodriguez-Esteban R, 2021). This citation distance differed from those described in previous work in that those typically involved directed connections (Botafogo et al., 1992). The citation distance between any pair of publications was computed using bidirectional breadth-first search (BFS) on citations existing at the time of publication of the latest article of the pair. Pairs of publications for which a path in the citation network could not be found were discarded from the analysis. A randomized version of the citation network was created by randomly swapping the nodes of the citation network, thus maintaining the network structure.

Citations came from the Open Citation Index repository (Peroni et al., 2017) and in particular from the March 23, 2020 update, which contained 721,655,465 citations between pairs of articles identified by a digital object identifier (DOI). DOI to PMID mappings were extracted from EBI’s PMID-PMCID-DOI dataset (Levchenko et al., 2018) downloaded on July 9, 2020, which contained 22,504,850 mappings between PMIDs and DOIs—thus covering 22,504,850 unique PMIDs in total. Using these mappings, 269,956,002 citations from the Open Citation Index were mapped from DOIs to PMIDs. As of July 2020, the fraction of publications covered by the Open Citation Index was 60% out of 51.1 million articles with references deposited with Crossref (https://i4oc.org/#about; checked on July 29, 2020).

The code used for this analysis is available at: https://github.com/raroes/scientific-silos

## 4. Results

Research on biomedical properties of compounds and genetic bases of disease is modeled here as a series of sequentially-built networks made of up to three nodes concerning each a gene, a compound and a disease. The nodes are connected by facts, which are molecular and medical relations published in the scientific literature. Central to this analysis is that two existing facts, e.g. a gene-disease and a disease-compound relation, precede and, therefore, *enable* the posterior new discovery of another fact, i.e. a gene-compound relation (Figure 1). For example, the compound isopropanol leads to increased expression of the gene NQO1’s mRNA (Vandebriel et al., 2010). This, together with the fact that inhibition of NQO1 is linked to the amelioration of kidney diseases (Chen et al., 2011), enables a new discovery, namely the relation between isopropanol and kidney diseases (Brott et al., 2013). Using a comprehensive dataset containing thousands of such facts, this model can be employed to understand the dynamics of scientific discovery.

**Figure 1.**
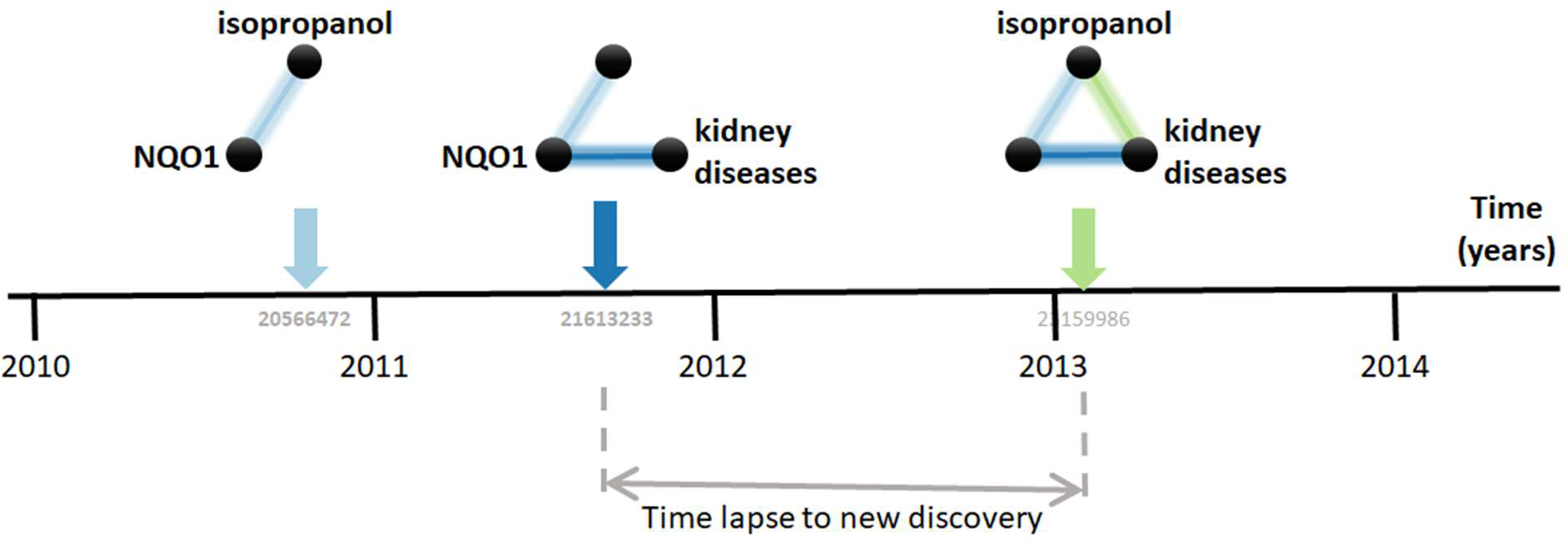
Connecting the dots. The three elements involved are the compound isopropanol, the gene NQO1 and kidney diseases. Each fact is a relation between two of these elements published in a MEDLINE article. For example, the relation between isopropanol and NQO1 was described in the article with PubMed ID 20566472 (Vandebriel et al., 2010). Data came from the Comparative Toxicogenomics Database (CTD).

The first step in the analysis is to find all combinatorially-possible pairs of facts sharing an element in the dataset, such as all pairs of facts involving the gene NQO1. Together, these facts comprise all pairs of facts that can enable new discoveries. If a pair of these facts is followed by a new discovery, the time lapsed until that event is computed. E.g., in Figure 1, the time lapse is between August 2011, when the second fact was published, and January 7, 2013, when the new discovery was published. This time lapse is then used to estimate the pace at which scientists produce new discoveries from existing facts and, in the case studied here, to test its dependence on the “distance” between the publications in which the facts were published. The distance metric used is the citation distance, which is a simple way to measure proximity in the scientific landscape (Rodriguez-Esteban, 2020). This distance is computed based on the citations existing at the time that the second fact is published. E.g., in Figure 1, the citation distance was 4 based on citation data from publications until August 2011, when the second fact was published.

The dataset used initially for this analysis was the Comparative Toxicogenomics Database (CTD) (Davis et al., 2019), which contains manually-curated relations between compounds, diseases and genes from the literature. Out of all combinatorially-possible pairs of facts in the CTD (n=6,261,706), only a small percentage (0.25%) was followed by new discoveries after 5 years. This percentage changed only slightly over the decades despite power-law growth in the combinatorial possibilities (Figure 2).

**Figure 2.**
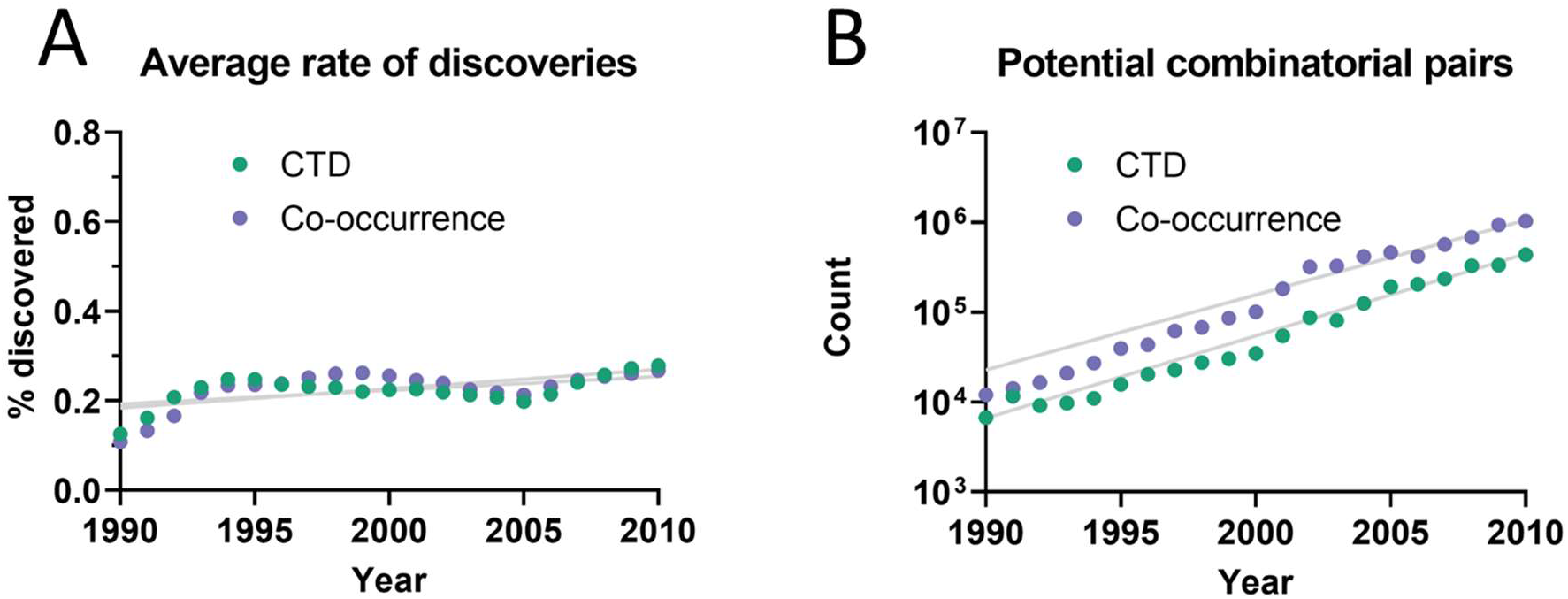
(A) Percentage of pairs of facts enabling discoveries after 5 years averaged over a 10-year time window. E.g., the earliest data point (1990) is an average for the period 1990-1999. Linear regressions were fitted to each curve. (B) Number of combinatorially-possible pairs of facts per year for each dataset. The year refers to the time when the second fact was published.

As can be seen in Figure 3A, the percentage of all combinatorially-possible pairs of facts that were followed by a new discovery increased linearly over the years, as scientists had time to work with them. This percentage, however, decreased with increasing citation distance, following an exponential decay (Figure 3B). For citation distance of 2, the percentage of combinatorially-possible pairs of facts enabling a new discovery was, on average, 0.090% per year, while for citation distance of 5 it was an average of 0.036%. After 5 years, it was 2.6 times more likely that a new discovery would be made out of facts separated originally by a citation distance of 2 than out of facts separated by a citation distance of 5 (0.47% vs. 0.18%).

**Figure 3.**
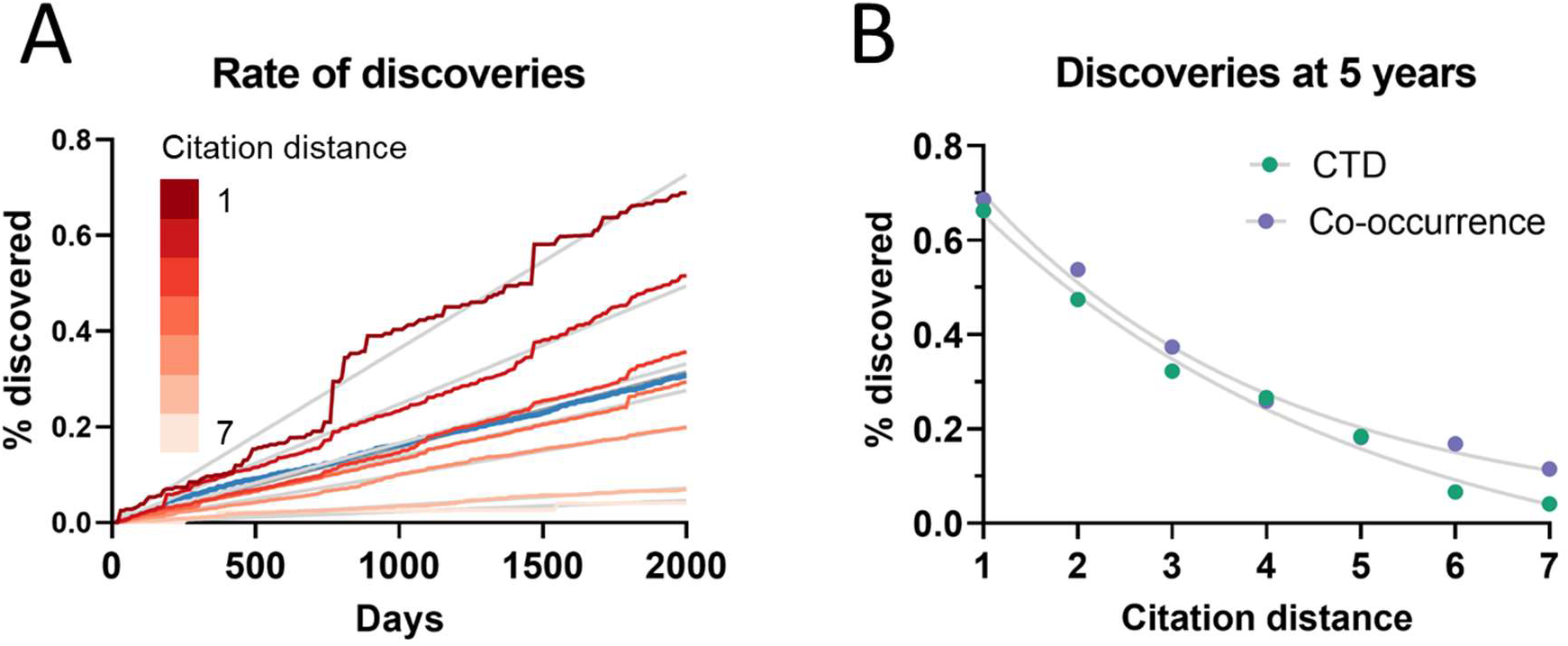
Percentage of pairs of facts enabling new discoveries (A) over time in CTD, and (B) after 5 years, based on citation distance. The percentage of pairs of facts enabling new discoveries increased faster over time with smaller citation distance. Origin-intercept linear (A) and exponential (B) regressions were fitted to each curve. The blue line in (A) represents the overall percentage trend for all distances.

This effect disappeared if all publications were randomly swapped within the citation network (Figure 4). In this case, the rate of discovery did not vary with citation distance, except for the case of distance equal to 1, due to data sparsity.

**Figure 4.**
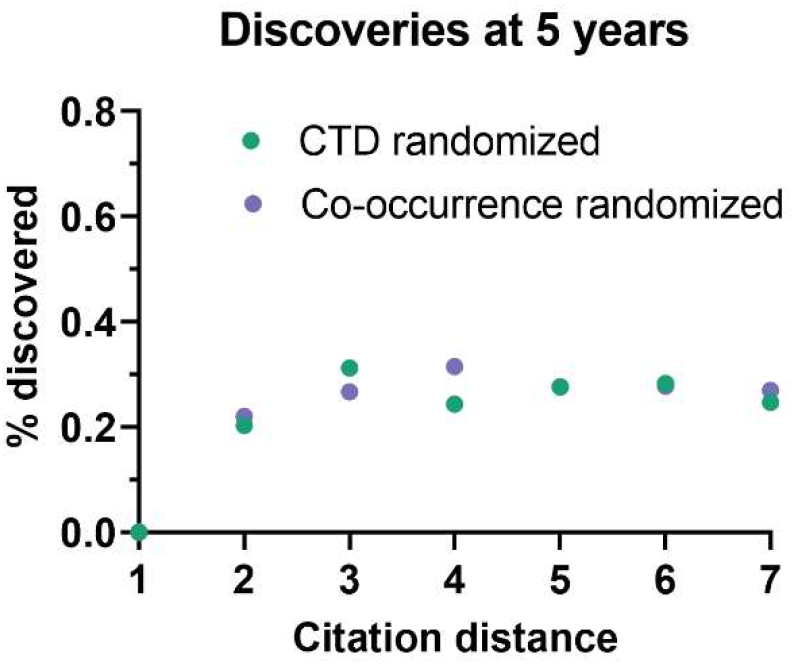
Percentage of pairs of facts enabling new discoveries after 5 years based on citation distance in a randomized citation network. Unlike in the non-randomized network, values did not follow an exponential decay based on citation distance.

To seek additional validation for these results, a similar analysis was performed with a different dataset based on co-occurrence of manual annotations of genes, diseases and chemicals/drugs of MEDLINE records. Co-occurrences have been considered suggestive of relations (Pavlopoulos et al., 2014) and have been used to discover new relations between drugs, genes and diseases (Frijters et al., 2010). The combinatorial space of all potentially enabling pairs of facts was three times larger (n=17,040,304) in this case than for CTD but the overall outcome was similar (Figure 5): Only a small percentage of those pairs of facts (0.26%) enabled new discoveries 5 years after publication. The percentage grew steadily with time, but at a different rate depending on the citation distance, following an exponential decay (Figure 3). For facts separated by a citation distance of 2, the percentage enabling a new discovery increased, on average, 0.10% per year, while for a citation distance of 5, it was 0.035%. After 5 years, it was 3 times more likely that a new discovery was be made out of facts published within a citation distance of 2 than out of facts within a citation distance of 5 (0.54% vs. 0.18%). This effect disappeared if publications were randomly swapped (Figure 4). Similarly to the previous case, the percentage of pairs that enabled a new discovery did not vary with citation distance and was similar to the baseline, except for distance equal to 1 due to data sparsity.

**Figure 5.**
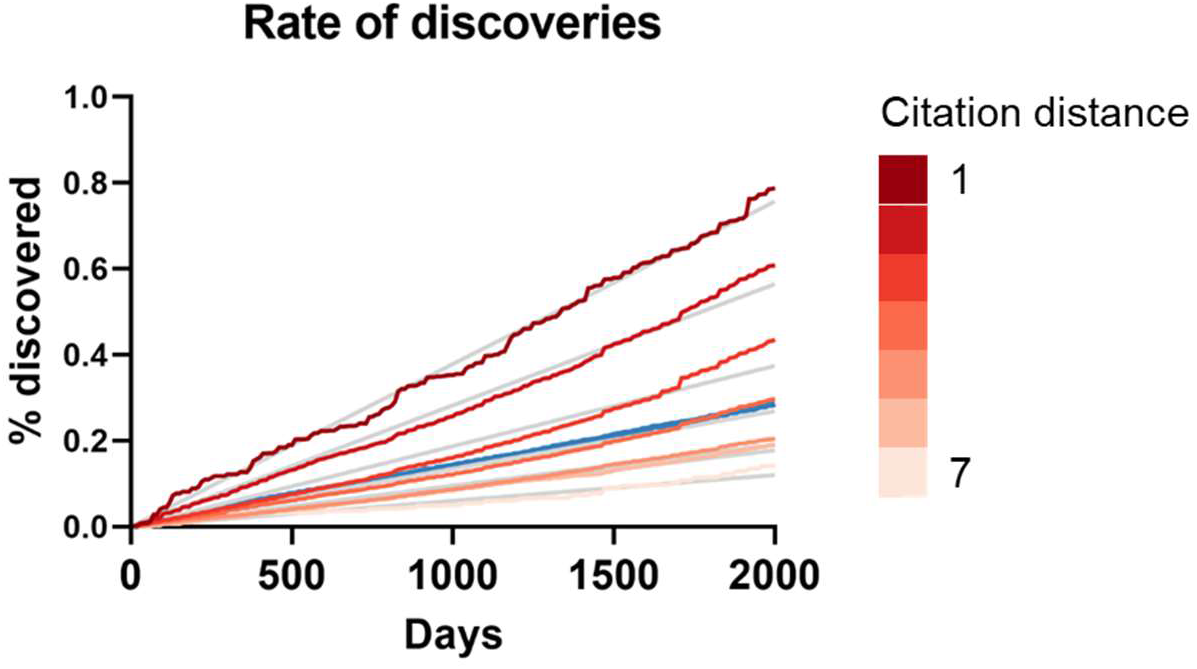
Percentage of pairs of facts enabling new discoveries over time based on citation distance in the co-occurrence dataset. Origin-intercept linear regressions were fitted to each curve. Quadratic regressions were a better fit. The blue line represents the percentage for all citation distances.

One potential weakness of this analysis could be missing citation data. The effect of this shortcoming was examined by eliminating existing citations randomly. This reduction did not change the shape of the outcome except when it was large (75% reduction) (Figure 6). Thus, an increase in the availability of citation data would not be expected to change the overall picture either.

**Figure 6.**
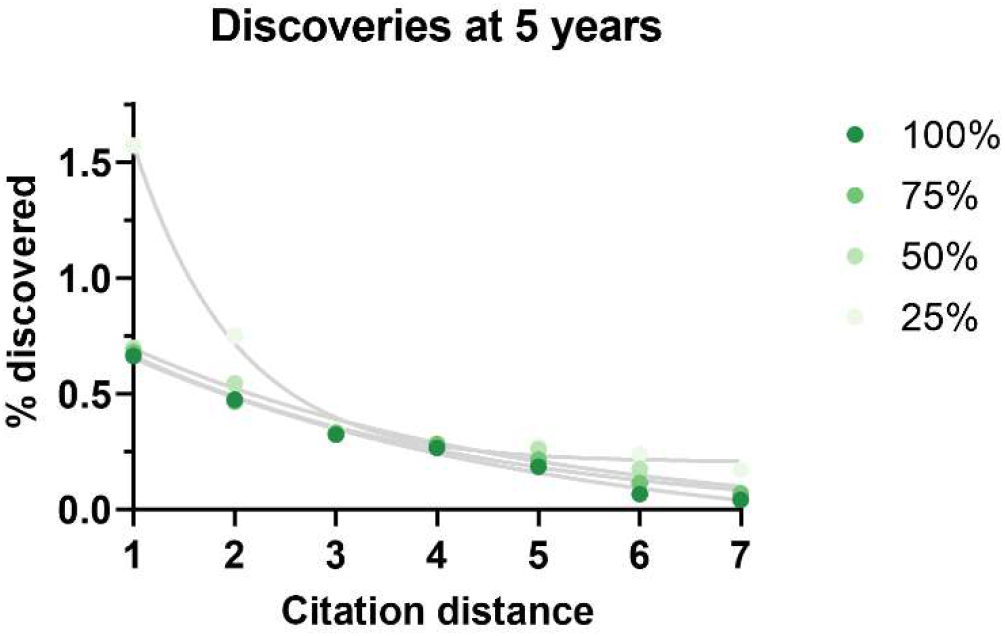
Percentage of pairs of facts enabling new discoveries after 5 years based on citation distance in a citation network with progressively less citations (100% = all citations available used, 75% = 75% of all citations available used, etc.). Exponential regressions were fitted to each curve. Data source was CTD.

## 5. Discussion and conclusions

The fact that the analyses on both datasets led to similar outcomes lends some validation to the results. Both analyses show that, over time, scientists “connect” only a small percentage of existing facts about relations between compounds, genes and diseases. Thus, biomedical scientists appear to have a wide set of facts available from which they only end up publishing discoveries about a small subset of them, whether because of lack of resources, lack of interest, or because many combinations lead to negative results. Moreover, scientists steadily “accumulate” discoveries over the years but the rate of collective accumulation is higher when those discoveries concern facts that were originally closer within the citation network. This points towards a path-dependency in scientific discovery (Rzhetsky et al., 2015) in which originally closely-located discoveries drive the sequence of future discoveries rather than optimal unbiased choices.

As more facts are discovered, one may expect their potential combinations to grow quadratically and siloization to be a consequence, at least partially, of this. However, there is a countervailing trend, which is that the scientific literature grows exponentially and it is able therefore to produce an increasingly larger number of discoveries. This analysis points to a somewhat stable relation between these two opposing forces. The overall percentage of facts that are being connected to form new discoveries has not changed much over the last decades and even increased slightly despite enormous growth in combinatorial possibilities (Figure 2). If scientists were falling behind, we would expect to see a decrease. Additionally, the rate of accumulation of new discoveries (Figures 3 and 5) appears generally stable and does not show signs of acceleration or deceleration over time (if only slight acceleration for co-occurrence data). Therefore, Swanson’s warning about “connection explosion” (Swanson, 2008) (“The literature of science cannot grow faster than the communities that produce it, but not so with connections. Implicit connections between subspecialties grow combinatorially. LBD is challenged more by a connection explosion than by an information explosion.”) does not bear on this case, probably because scientists tend to lend a higher focus to a reduced set of drugs, diseases and genes (Yao et al., 2015; Haynes et al., 2018; Stoeger et al., 2018; Rzhetsky et al., 2015), which would tend to limit combinatorial explosion.

The citation distance is only a rough estimate of scientific proximity between articles. One could expect that a more precise surrogate for scientific proximity could show an even stronger siloization effect. The citation distance was chosen for its simplicity. Measures of semantic similarity between articles, for example, could create cross-feedback between article annotations (i.e. gene annotations) and the distance metric itself.

Reaching more often for facts that are “closer” could be a simple heuristic or a type of availability bias. That scientists may use heuristic biases, even if unconscious, to select their research goals should not be surprising, given the extraordinary growth of the scientific literature in most fields. However, this bias leads to a distortion of scientific progress and an opportunity for those who may venture further away from their silos with the aid of modern tools (Krenn & Zeilinger, 2020; Whalen et al., 2016). Siloization is ultimately an emerging property of scientific organization and self-organization with cognitive, social and technological aspects.

## Declarations

### Availability of data and material

All data used in this study is publicly available.

### Code availability

The code used for this analysis is available at: https://github.com/raroes/scientific-silos

## References

Auffray C, Chen Z, Hood L. Systems medicine: the future of medical genomics and healthcare. Genome Med. 2009 Jan 20;1(1):2.

Baumwol K, Mortimer ST, Huerta TR, Norman CD, Buchan AMJ. Promoting interdisciplinarity in the life sciences: a case study. Res Eval. 2011;20(4):283–292.

Bekhuis T. Conceptual biology, hypothesis discovery, and text mining: Swanson’s legacy Biomed Digit Libr. 2006;3.

Bornmann L, Mutz R. Growth rates of modern science: A bibliometric analysis based on the number of publications and cited references. J Assn Inf Sci Tec, 66: 2215–2222. 2015.

Botafogo RA, Rivlin E, Shneiderman B. Structural analysis of hypertexts: Identifying hierarchies and useful metrics. ACM Transactions on Information Systems. 1992;10(2):142–180.

Bromham L, Dinnage R, Hua X. Interdisciplinary research has consistently lower funding success. Nature. 2016 Jun 30;534(7609):684–7.

Brott DA, Bentley P, Nadella MV, Thurman D, Fikes J, Cheatham L, McGrath F, Luo W, Kinter LB. Renal biomarker changes associated with hyaline droplet nephropathy in rats are time and potentially compound dependent. Toxicology. 2013 Jan 7;303:133–8.

Bruggeman J, Traag VA, Uitermark J. Detecting Communities through Network Data. Am Soc Rev. 2012;77(6):1050–1063.

Chen M, Gong L, Qi X, Xing G, Luan Y, Wu Y, Xiao Y, Yao J, Li Y, Xue X, Pan G, Ren J. Inhibition of renal NQO1 activity by dicoumarol suppresses nitroreduction of aristolochic acid I and attenuates its nephrotoxicity. Toxicol Sci. 2011 Aug;122(2):288–96.

Cokol M, Iossifov I, Weinreb C, Rzhetsky A. Emergent behavior of growing knowledge about molecular interactions. Nat Biotechnol. 2005 Oct;23(10):1243–7.

Davis AP, Grondin CJ, Johnson RJ, Sciaky D, McMorran R, Wiegers J, Wiegers TC, Mattingly CJ. The Comparative Toxicogenomics Database: update 2019. Nucleic Acids Res. 2019 Jan 8;47(D1):D948–D954.

Fortunato S, Bergstrom CT, Börner K, Evans JA, Helbing D, Milojević S, Petersen AM, Radicchi F, Sinatra R, Uzzi B, Vespignani A, Waltman L, Wang D, Barabási AL. Science of science. Science. 2018 Mar 2;359(6379):eaao0185.

Frijters R, van Vugt M, Smeets R, van Schaik R, de Vlieg J, Alkema W. Literature mining for the discovery of hidden connections between drugs, genes and diseases. PLoS Comput Biol. 2010 Sep 23;6(9):e1000943.

Haynes WA, Tomczak A, Khatri P. Gene annotation bias impedes biomedical research. Sci Rep. 2018 Jan 22;8(1):1362.

Heimeriks G, Boschma R. The path- and place-dependent nature of scientific knowledge production in biotech 1986–2008. J Econ Geogr. 2014;14(2):339–64.

Krenn M, Zeilinger A. Predicting research trends with semantic and neural networks with an application in quantum physics. Proc Natl Acad Sci U S A. 2020 Jan 28;117(4):1910–1916.

Larsen PO, von Ins M. The rate of growth in scientific publication and the decline in coverage provided by Science Citation Index. Scientometrics. 2010. 84(3): 575–603.

Leischow SJ, Best A, Trochim WM, Clark PI, Gallagher RS, Marcus SE, Matthews E. Systems thinking to improve the public’s health, Am J Prev Med. 2008 Aug;35(2 Suppl):S196–203.

Levchenko M, Gou Y, Graef F, Hamelers A, Huang Z, Ide-Smith M, Iyer A, Kilian O, Katuri J, Kim JH, Marinos N, Nambiar R, Parkin M, Pi X, Rogers F, Talo F, Vartak V, Venkatesan A, McEntyre J. Europe PMC in 2017. Nucleic Acids Res. 2018 Jan 4;46(D1):D1254–D1260.

Luke DA, Carothers BJ, Dhand A, Bell RA, Moreland-Russell S, Sarli CC, Evanoff BA. Breaking down silos: mapping growth of cross-disciplinary collaboration in a translational science initiative. Clin Transl Sci. 2015 Apr;8(2):143–9.

Maglott D, Ostell J, Pruitt KD, Tatusova T. Entrez Gene: gene-centered information at NCBI. Nucleic Acids Res. 2011 Jan;39(Database issue):D52–7.

Pavlopoulos GA, Promponas VJ, Ouzounis CA, Iliopoulos I. Biological information extraction and co-occurrence analysis. Methods Mol Biol. 2014;1159:77–92.

Peroni S., Shotton D., Vitali F. One Year of the OpenCitations Corpus. In: d’Amato C. et al. (eds) The Semantic Web – ISWC 2017. ISWC 2017. Lecture Notes in Computer Science 2017; 10588. Springer, Cham.

Rodriguez-Esteban R, Loging WT. Quantifying the complexity of medical research. Bioinformatics. 2013 Nov 15;29(22):2918–24.

Rodriguez-Esteban R. Semantic persistence of ambiguous biomedical names in the citation network. Bioinformatics. 2020 Apr 1;36(7):2224–2228.

Rodriguez-Esteban R. Biomedical articles share annotations with their citation neighbors. BMC Bioinformatics. 2021 Feb 26;22(1):95.

Rzhetsky A, Foster JG, Foster IT, Evans JA. Choosing experiments to accelerate collective discovery. Proc Natl Acad Sci U S A. 2015 Nov 24;112(47):14569–74.

Shia F, Foster JG, Evans JA. Weaving the fabric of science: Dynamic network models of science’s unfolding structure. Soc Netw. 2015. 43:73–85.

Sieloff CG. “If only HP knew what HP knows”: the roots of knowledge management at Hewlett-Packard. J Knowledge Management. 1999;3(1):47–53.

Smalheiser NR. Literature-based discovery: Beyond the ABCs. J Am Soc Inf Sci Tech. 2012;63(2):218–24

Soler L, Trizio E, Pickering A. Science as It Could Have Been: Discussing the Contingency/Inevitability Problem. University of Pittsburgh Press; 2015.

Stoeger T, Gerlach M, Morimoto RI, Nunes Amaral LA. Large-scale investigation of the reasons why potentially important genes are ignored. PLoS Biol. 2018 Sep 18;16(9):e2006643.

Swanson D R. Undiscovered public knowledge. Libr Q. 1986;56:103–118.

Törmä P. Scientific silos are holding back collaboration and breakthroughs. The Engineer. 2019 Nov 28.

Swanson DR. Literature-Based Discovery? The Very Idea. In: Bruza P., Weeber M. (eds) Literature-based Discovery. Information Science and Knowledge Management, vol 15. Springer, Berlin, Heidelberg. 2008.

Tambolo L. Counterfactual Histories of Science and the Contingency Thesis. In: Magnani L., Casadio C., eds. Model-Based Reasoning in Science and Technology. Studies in Applied Philosophy, Epistemology and Rational Ethics. Springer, Cham; 2017.

Thilakaratne M, Falkner K, Atapattu T. A Systematic Review on Literature-based Discovery: General Overview, Methodology, & Statistical Analysis. ACM Computing Surveys. 2019:129.

Vandebriel RJ, Pennings JL, Baken KA, Pronk TE, Boorsma A, Gottschalk R, Van Loveren H. Keratinocyte gene expression profiles discriminate sensitizing and irritating compounds. Toxicol Sci. 2010 Sep;117(1):81–9.

Vodovotz Y., An G. An Overview of the Translational Dilemma and the Need for Translational Systems Biology of Inflammation. In: Vodovotz Y., An G. (eds) Complex Systems and Computational Biology Approaches to Acute Inflammation. Springer, New York, NY. 2013.

Whalen R, Huang Y, Tanis C, Sawant A, Uzzi B, Contractor N. Citation Distance: Measuring Changes in Scientific Search Strategies. In Proceedings of the 25th International Conference Companion on World Wide Web (WWW ‘16 Companion). International World Wide Web Conferences Steering Committee, Geneva, Switzerland, 419–423. 2016

Yao L, Li Y, Ghosh S, Evans JA, Rzhetsky A. Health ROI as a measure of misalignment of biomedical needs and resources. Nat Biotechnol. 2015 Aug;33(8):807–11.

